# Order Matters: Ambient RNA correction confounds doublet detection in single-cell RNA sequencing

**DOI:** 10.64898/2026.10.13.758369

**Authors:** Michelle Meier, George Howitt, Calandra V Grima, Gunjan Dixit, Hamish King, Liam Gubbels, Eric Levi, Karen Davies, Shivanthan Shanthikumar, Melanie R. Neeland, Jovana Maksimovic, Alicia Oshlack

**Affiliations:** Peter MacCallum Cancer Centre, Melbourne, Victoria 3000, Australia; Sir Peter MacCallum Department of Oncology, The University of Melbourne, Victoria 3010, Australia; Walter and Eliza Hall Institute, Parkville, VIC, Australia; Infection, Immunity and Global Health, Murdoch Children’s Research Institute, Parkville, Australia; Department of Otolaryngology, Royal Children’s Hospital, Parkville, Australia; Clinical Sciences, Murdoch Children’s Research Institute, Parkville, Australia; Respiratory Diseases, Murdoch Children’s Research Institute, Parkville, VIC, Australia; Respiratory and Sleep Medicine, Royal Children’s Hospital, Parkville, VIC, Australia; Department of Paediatrics, University of Melbourne, Parkville, VIC, Australia; School of Mathematics and Statistics, The University of Melbourne, Victoria 3010, Australia

**Keywords:** single-cell RNA-sequencing, doublets, ambient RNA correction, workflows

## Abstract

A common technical artifact in droplet-based single-cell RNA sequencing (scRNA-seq) is the capture of two cells in the same droplet, termed doublets. Doublets may present as hybrid cell types, and their removal is a critical step during quality control. Another form of contamination is ambient RNA, which originates from free floating RNA released from other cells in suspension, likely during cell death. As with doublets, ambient RNA results in unexpected gene expression patterns. Most suggested analysis pipelines include ambient RNA correction before doublet detection however no formal investigation into how doublet detection is affected by ambient RNA correction has been conducted.

Here, we evaluate the impact of ambient RNA correction on the performance of doublet detection using novel engineered and multiplexed datasets. We apply three of the most widely-used methods for removing ambient RNA (*decontX*, *soupX*, *CellBender*), in conjunction with up to six doublet detection methods (*scDblFinder*, *scds*, *bcds*, *cxds, scrublet*, *DoubletDetection*). We find that doublet detection tools perform best when done prior to ambient RNA correction, with the exception of *CellBender*. Performing ambient RNA correction first, particularly for *decontX*, leads to an increased proportion of doublet droplets retained in the dataset compared with calling doublets on uncorrected data. We show that both *decontX* and *soupX* partially mask doublet signal by removing proportionally more ambient RNA from doublets that remain undetected after ambient RNA correction. Our results show that the performance of doublet detection is significantly impacted by the use of ambient RNA correction tools. We suggest a new workflow that implements doublet detection prior to ambient RNA correction unless using *CellBender*.

## Introduction

Single-cell RNA sequencing (scRNA-seq) has played a pivotal role in studying the expression profiles of thousands of individual cells [1,2]. Since its introduction, scRNA-seq has had a profound impact on the identification and characterisation of cell types and cell states [3] and our understanding of cellular heterogeneity [4,5]. Droplet-based scRNA-seq protocols have become particularly popular due to their high-throughput and low cost per cell [1,2,6]. Ideally, each cell is partitioned into a droplet containing an oligo barcoded bead which hybridises to the RNA molecules in the cell. However, a common technical artefact of droplet-based technologies is the capture of two or more cells in the same droplet, creating so-called “doublets”, whose pooled RNA expression is assigned to the same barcode [1]. The fraction of doublets is not insignificant and is directly proportional to the number of cells being sequenced. Typically, scRNA-seq experiments contain approximately 10-20% doublets [7].

As scRNA-seq experiments continue to grow in scale, multiplexing technology allows for cells from different samples to be pooled in a single capture [8]. The sample of origin for each cell can be tracked using either oligo-tagged antibodies, lipid-modified oligos [8,9] or natural genotypic diversity [10,11]. While multiplexing reduces both batch effects and sequencing costs, it often requires an increased capture size, leading to an increased proportion of doublets. Cell tagging allows demultiplexing tools to detect doublets formed by cells from different samples, and mark them for downstream removal [12].

A recent addition to the droplet-based scRNA-seq technology repertoire is the Flex protocol from 10x Genomics; a probe-based method for gene expression quantification that natively supports multiplexed experimental designs [13]. Each sample is mixed with sample barcoded gene probes that hybridise to the RNA molecules in suspension, tagging each sequence/oligo with a sample-specific barcode. Samples are then pooled, captured and sequenced using canonical droplet-based technology. Consequently, each read can be traced back to its sample of origin, as well as the droplet it was captured in. This allows both the identification of between-sample doublets, as well as their resolution back into single cells. However, demultiplexing still fails to identify doublets formed by cells from the same sample, meaning doublet detection based on transcriptomic signals remains a necessary processing step [14,15].

Broadly, doublets can be split into two main classes [16,17]: homotypic doublets capture two cells of the same, or transcriptionally very similar cell types, while heterotypic doublets capture two cells with distinct expression profiles. Heterotypic doublets, in particular, can negatively impact downstream analysis as they often adopt a hybrid cell type identity which can lead to the misidentification of novel cell states [18]. In addition, it has been shown that doublets can impact clustering, differential gene expression analysis and inference of developmental trajectories [15,19]. It is therefore critical to accurately identify and remove most doublets prior to downstream analysis [18,20,21].

Numerous different methods have been developed to identify and remove doublets in scRNAseq datasets [19]. The majority of these tools generate artificial doublets by combining gene expression profiles of two droplets in the dataset [7,14,16,17,22–27]. Real doublets are then identified by evaluating their similarity to artificial doublets. The exception to this approach is the *cxds* tool, which uses the co-expression of mutually exclusive marker genes to identify doublets [24]. While benchmarking studies evaluating and comparing the performance of these tools highlight that there is no universally best tool [19,28], *scDblFinder* tends to outperform other methods in terms of accuracy and its ability to improve downstream analysis [29].

Another technical artefact of droplet-based sequencing is the contamination of droplets by cell-free RNA molecules, called ambient RNA. These exogenous RNA molecules that are captured in the droplet alongside a cell, are assigned to the same barcoded bead, and can contribute 3-35% of the droplet’s total reads [30]. The amount of contaminating ambient RNA in a dataset can be highly variable and difficult to predict. However, even high-quality datasets can contain significant ambient RNA [31] and it tends to be more abundant in complex biological settings, such as the brain [32]. In addition, ambient RNA is more common in single-nuclei RNA-seq samples prepared from fresh-frozen tissue [30,33]. Importantly, ambient RNA has been shown to impact cell clustering, the detection of marker genes, creates batch effects and can even lead to misannotation of cell types [30,32–34].

For these reasons, it is important to quantify and remove ambient RNA expression. Several tools have been developed to remove ambient RNA contamination across each droplet in an experiment [33]. Multiple independent benchmarks have identified *CellBender* [34], *soupX* [31] and *decontX* [35] as the most reliable ambient RNA correction tools [30,33]. *CellBender* tends to provide the best performance, but its application is severely limited by the computational resources it requires even for a low number of cells. Performance for *soupX* and *decontX* varies across datasets but offer satisfactory computationally lightweight alternatives. In contrast to *CellBender* and *soupX*, *decontX* does not require empty droplets to perform ambient RNA correction, making it a popular choice for re-analysis of public data, where empty droplets may not be included. All three of these ambient RNA correction tools are either designed or suggested to be run upstream of any other form of quality control, including doublet detection. Consequently, ambient RNA correction is performed prior to doublet detection in current standardised scRNA-seq processing workflows [20,36,37].

Ambient RNA contamination and doublets are conceptually similar forms of contamination [19]. Both pollute the droplet’s content with RNA molecules that are exogenous to the captured cell being profiled. This is exemplified by Yang *et al*. [35], who found that predicted doublets exhibit a much higher proportion of estimated contamination than singlets. In addition, the impact of both ambient RNA and doublets on downstream analysis are similar, both being shown to affect clustering, reduce effect size in marker gene analysis and lead to misannotations of cell types [19,30].

Despite these evident similarities between ambient RNA contamination and doublets, there has been limited investigation into how doublet detection and ambient correction tools interact. The authors of *CellBender* demonstrate that its ambient correction performance is unaffected by the presence of doublets and Neavin *et al.* [14] show that most doublet detection tools are robust to moderate levels of ambient RNA. Doublets and ambient RNA contamination are handled through fundamentally different strategies: ambient RNA correction generates an altered (ambient corrected) count matrix which is used for all downstream analysis (including doublet detection), whereas doublets are typically removed from the analysis entirely.

In this study, we assess how ambient RNA correction affects the performance of doublet detection. Using different single-cell technologies across different biological conditions, we apply three of the most widely-used ambient RNA correction methods (*decontX*, *soupX*, *CellBender*) in conjunction with up to six doublet detection methods (*scDblFinder*, *scds*, *bcds*, *cxds*, *scrublet*, *DoubletDetection*). Overall, our results highlight that doublet detection is affected by the use of the ambient RNA correction tools. We conclude that in many settings doublet detection should be performed prior to ambient RNA correction and propose a guide to determine the order that the tools tested in this study should be implemented in future workflows.

## Results

### Performing ambient RNA correction alters doublet detection outcomes

To investigate whether ambient RNA correction affects doublet calling, we use three different datasets: a 16-plex Flex dataset of healthy peripheral blood mononuclear cells (PBMCs) provided by one donor (the PBMC dataset), a 16-plex Flex scRNA-seq dataset consisting of tonsillar tissue derived from 16 healthy paediatric donors (the tonsil dataset), and a 2-plex Flex dataset of two ovaries donated by one ovarian cancer patient (the ovarian cancer dataset). These datasets represent a spectrum of biological and technical complexity, ranging from healthy with few expected immune interactions (PBMC dataset), to healthy with high immune activity (tonsil dataset) to complex disease (ovarian cancer dataset). We remove ambient RNA contamination using *decontX* and identify doublets in both the ambient corrected and uncorrected datasets using *scDblFinder* (Figure 1A). We selected *decontX* due to its ability to perform ambient RNA correction without empty droplets, and *scDblFinder* due to its noted high performance in multiple independent benchmarks. In the absence of ground truth labels, we define the uncorrected workflow as a baseline against which the effects of ambient RNA correction are assessed.

**Figure 1.**
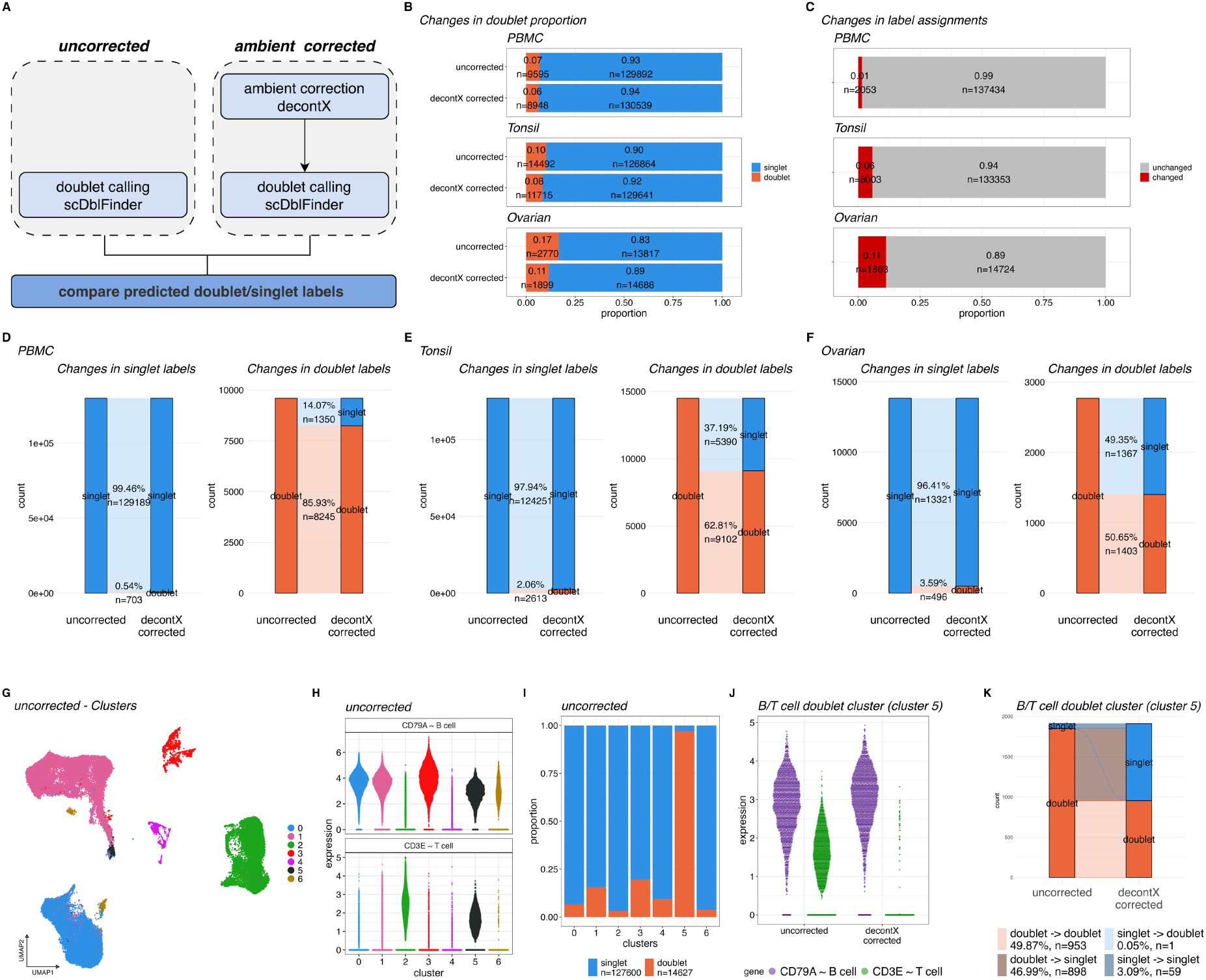
Performing ambient RNA correction alters doublet detection outcomes. **A.** Overview of workflow to compare predicted doublet labels before and after ambient RNA correction using decontX for all datasets. Ambient RNA correction is performed using decontX, doublet detection is performed using scDblFinder. **B.** Stacked barplots showing the proportion of doublets and singlets in the uncorrected (top) and decontX corrected (bottom) datasets. **C.** Stacked barplots showing the proportion of cells whose predicted labels change after ambient correction. **D-F** River plots showing how predicted labels change for singlets (left) and doublets (right) separately for the PBMC (**D**), tonsil (**E**) and ovarian cancer (**F**) dataset. **G.** UMAP visualisation showing clusters in the uncorrected tonsil dataset. **H**. Violin plot showing gene expression for CD79A (B cell marker) and CD3E (T cell marker) for the clusters in the uncorrected dataset. **I.** Stacked barplots showing the proportion of doublets and singlets for each cluster in the uncorrected dataset. **J.** Violin plot showing gene expression changes for CD79A (B cell marker) and CD3E (T cell marker) in B/T cell doublet cluster (cluster 5) cells (black, n=1911) before (left) and after (right) ambient RNA correction. **K.** River plot showing how doublet labels change for uncorrected doublet cluster cells before (left) and after (right) ambient RNA correction for B/T cell doublet cluster (cluster 5) cells.

We first compare the number of doublets detected with and without ambient correction (Figure 1B). For the PBMC dataset the overall proportion of doublets after ambient RNA correction is comparable to the uncorrected workflow, while the proportion of doublets in the corrected workflow is slightly lower for the tonsil dataset. In the ovarian cancer dataset, the decrease in the proportion of doublets is more pronounced, dropping from 0.17 to 0.11 after ambient RNA correction (Figure 1B). While most cells have the same labels before and after ambient correction in the PBMC dataset, the labels change for around 8,000 cells (6% of dataset) in the tonsil dataset (Figure 1C). The proportion of changed labels is even greater in the ovarian cancer dataset (11% of dataset), which aligns with the more significant decrease in doublets observed in this dataset (Figure 1B). Finally, we explore the proportion of cells whose labels change for singlets and doublets separately (Figure 1D-F). Generally, most singlets do not change their labels (>96% unchanged). The proportion of doublets whose labels change to singlet is low in the PBMC dataset (Figure 1D) and increases in the tonsil (Figure 1E) and to an even greater extent in the ovarian cancer dataset (Figure 1F). Notably, almost 50% of doublets change their labels to singlets in the ovarian cancer dataset. Similar results are seen in non-flex datasets when running the same workflow on a 10x 3’ v2 and 10x 5’ v1 tonsil sample (Supplementary Figure 1A-H).

Together this shows that removing ambient RNA prior to doublet detection alters the outcome of doublet detection. Moreover, the discordance of doublet labels before and after ambient correction appears to be correlated with biological complexity.

To further investigate the impact of changes to doublet labels, we perform basic cell clustering and labelling of the tonsil dataset using the workflow with or without *decontX* ambient correction. For both the uncorrected and corrected datasets, we identify clusters of CD79A^+^ B cells, CD3E^+^ T cells, and CD79A^-^ CD3E^-^ non-lymphoid cells (Figure 1G-H). It is common for heterotypic doublets to form their own clusters, which gives them the appearance of hybrid, transitional or novel cell types [17]. Indeed, in the uncorrected dataset, we observe putative doublets enriched in cluster 5, whose cells co-express B and T cell markers (Figure 1H-I). Interestingly, such a doublet cluster is absent in the ambient corrected dataset (Supplementary Figure 1I-K). Upon further inspection of the cells in cluster 5 (B/T cell doublet cluster), we see a dramatic reduction of *CD3E* expression in these putative doublets post ambient RNA correction (Figure 1J), giving them a B-cell singlet-like appearance. In addition, most (97.8%, n=1869) of these cells are assigned to CD79A^+^ B cell clusters after ambient RNA correction (Supplementary Figure 1L). Over 45% (n=898) of these cells change their predicted label from doublet to singlet after ambient RNA correction (Figure 1K). This suggests that ambient correction removes the T-cell signature from one cell in the doublet making the droplet resemble a singlet to the doublet caller.

### Generating engineered datasets from 10x Flex data to evaluate doublet composition and detection performance

Benchmarking studies for doublet detection tools commonly use multiplexed data with ground truth doublet labels generated from droplets containing two samples. However, these datasets cannot provide insight into the cell type composition of doublets. The ability to characterise doublets in-depth, including their composition, may help to explain why some doublets appear to be more sensitive to ambient RNA correction than others. Therefore, we utilise the multiplexed 10x Flex data to generate datasets with known doublet labels and composition.

The 10x Flex protocol is a probe-based method for gene expression quantification. Each read carries both a sample as well as a droplet barcode (Figure 2A). This allows every read to be demultiplexed into the samples of origin, allowing deconvolution of multiple cells in each droplet. The combination of all cells carrying the same droplet barcode are referred to as *droplets*, while we refer to unique sample and droplet barcode combinations as *pseudo-droplets.* Pseudo-droplets are the outputs of the Cell Ranger pipeline for Flex data, and are the equivalent to droplets in canonical single-cell experiments. To make a clean dataset, we remove same-sample doublets and low quality pseudo-droplets before combining pseudo-droplets that carry the same droplet barcode to form engineered droplets. This generates singlets, as well as engineered doublets as they would have occurred during the droplet capture (Figure 2B). We also collect metadata information for individual cells in each engineered droplet (see Methods). This allows us to further annotate and describe the cells that make up a doublet.

**Figure 2.**
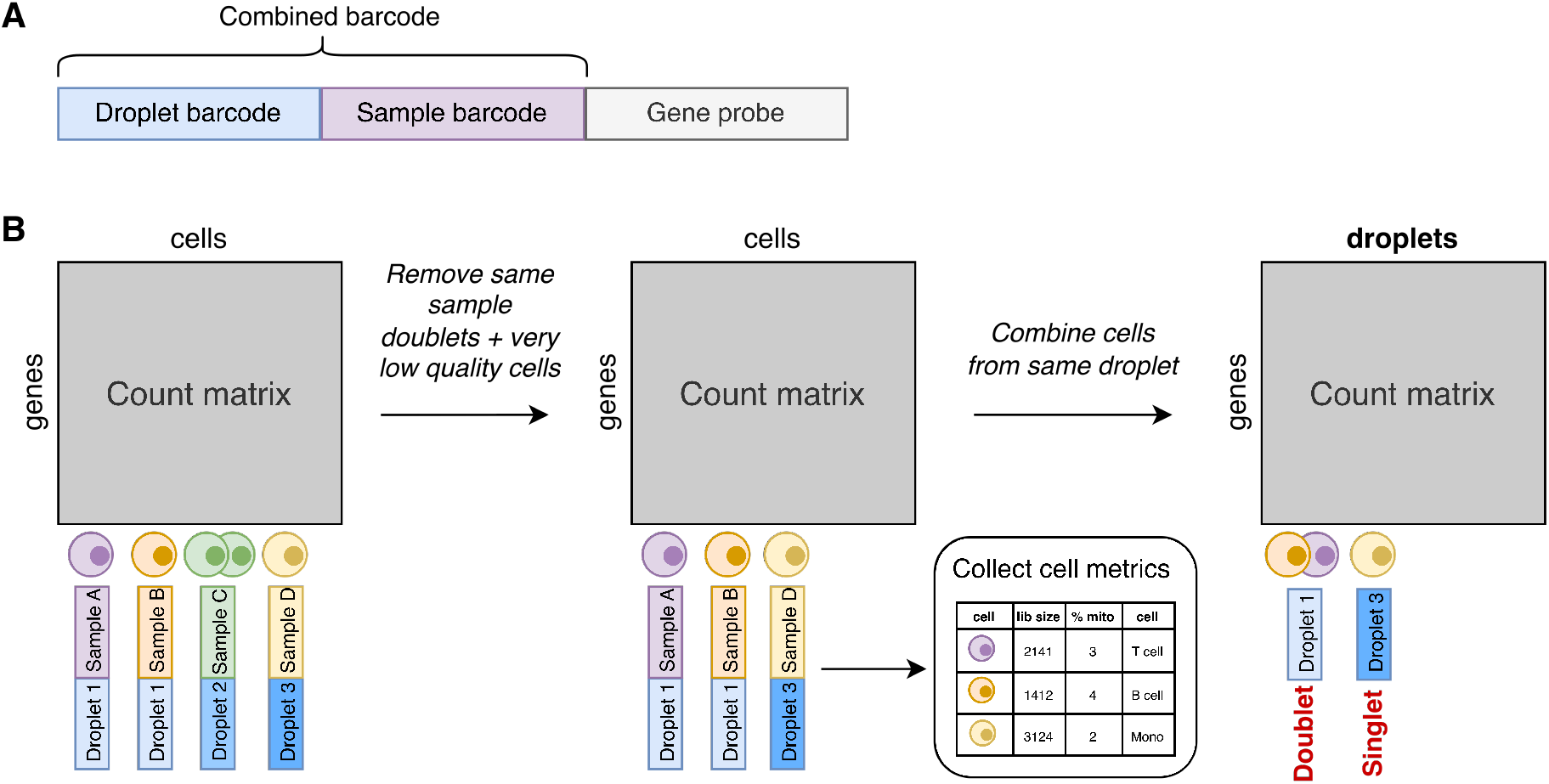
Generating engineered datasets from 10x Flex data to evaluate doublet composition and detection performance. **A.** Simplified illustration of 10x Flex barcode structure. The first section of the barcode represents the droplet barcode, followed by the sample barcode. **B.** Overview of workflow to simulate datasets using 10x Flex data.

Using the tonsil dataset, we generate a dataset with ground truth doublets using the engineered droplets. To appropriately mimic real-life dataset metrics, we sampled the dataset to 12,000 droplets with a doublet proportion of 0.15 (10,200 singlets + 1,800 doublets). We refer to this dataset as the engineered tonsil dataset. In total, we generate 5 batches of engineered datasets by repeating the subsampling step.

### Doublet detection performance is reduced after ambient RNA correction in engineered datasets

Using the engineered tonsil datasets, we investigate how ambient RNA correction affects the performance of doublet detection. We apply three of the most widely-used methods for removing ambient RNA: *decontX*, *soupX* and *CellBender* (Figure 3A). For *decontX*, we employ both settings either with or without providing empty droplets as background (decontX_BG and decontX respectively). Doublets are identified using *scDblFinder* before and after ambient RNA correction. We perform this evaluation for all 5 engineered dataset batches. Settings and parameters for all ambient correction methods and *scDblFinder* are included in the Methods section.

**Figure 3:**
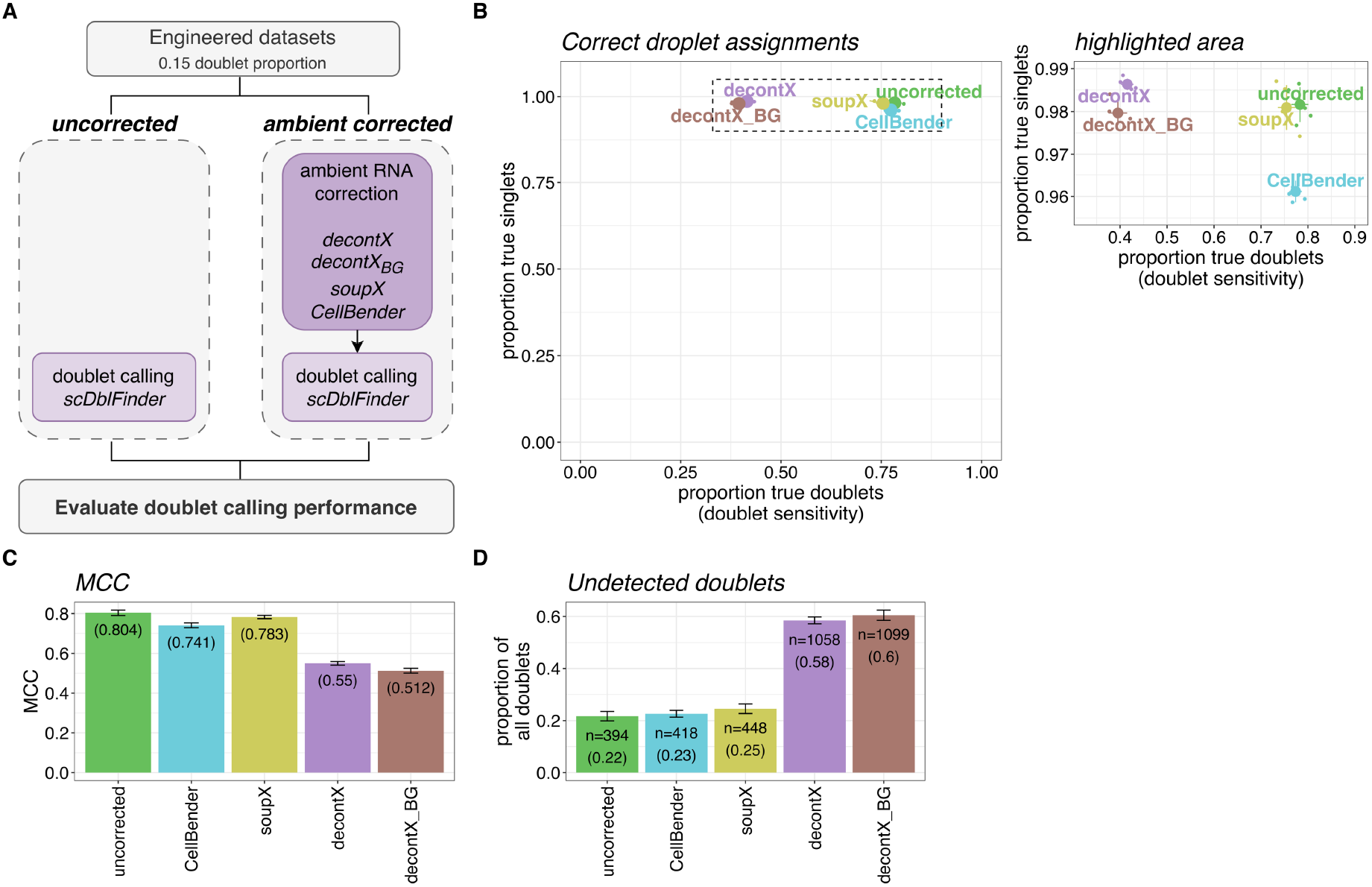
Doublet detection performance is reduced after ambient RNA correction in engineered datasets. **A.** Schematic overview of workflow to compare doublet detection performance in uncorrected and ambient RNA corrected settings. Ambient RNA correction is performed using soupX, decontX, decontX with background (decontX_BG) or CellBender. Doublet calling is performed using scDblFinder. **B.** Scatterplot showing proportion of true singlets (y-axis) and true doublets (x-axis, doublet sensitivity) detected for all batches (small dots), as well as mean across all batches (large dots). Lines extend to +/-1 standard deviation across batches. **C-D.** Barplots showing the mean MCC (**C**) and mean proportion of undetected doublets (**D**) for each ambient RNA correction setting. The number of undetected doublets is also provided in **D**. Whiskers extend to +/-1 standard deviation across batches.

To assess the doublet calling performance for all configurations we quantify both the proportion of true singlets and doublets that are assigned correctly, as well as the Matthews correlation coefficient (MCC). Overall, we observe that the proportion of true singlets detected is consistently high (mean 0.978), while the proportion of true doublets detected (doublet sensitivity) varies greatly (0.395-0.783) (Figure 3B). We find that, on average, the highest doublet sensitivity is achieved when doublet detection is performed on data without ambient RNA correction (Figure 3B, Supplementary Figure 2A). This coincides with an increased MCC for doublet calling in the uncorrected dataset compared to all ambient RNA corrected datasets, indicating an improved, balanced doublet calling performance in general (Figure 3C, Supplementary Figure 2B, Extended Data Table 01). Particularly for *decontX*, ambient RNA correction leads to an increased proportion of undetected doublets which remain in the dataset as singlets (Figure 3D). Critically, these mislabelled doublets would not be removed during quality control, which may cause unwanted retention of doublets for downstream analysis.

### Engineered doublets reveal key features of undetected doublets after ambient RNA correction

We next investigate doublets that are undetected in different settings to further understand what may cause a doublet to evade detection following different ambient RNA correction tools. To avoid double counting, we remove droplets that randomly appear in multiple batches, leaving a total of 5,172 doublets and 5,085 singlets (Supplementary Figure 3). Across all settings, we find n=1,663 doublets that are always detected and n=809 doublets that are never detected in any setting (Figure 4A, orange and grey, respectively). The “never detected” set of doublets serves as a control for the inherent limitations of *scDblFinder*. We compare the remaining doublets that were not detected after each ambient RNA correction setting (Figure 4A, dark shading) to those that are always or never detected.

**Figure 4.**
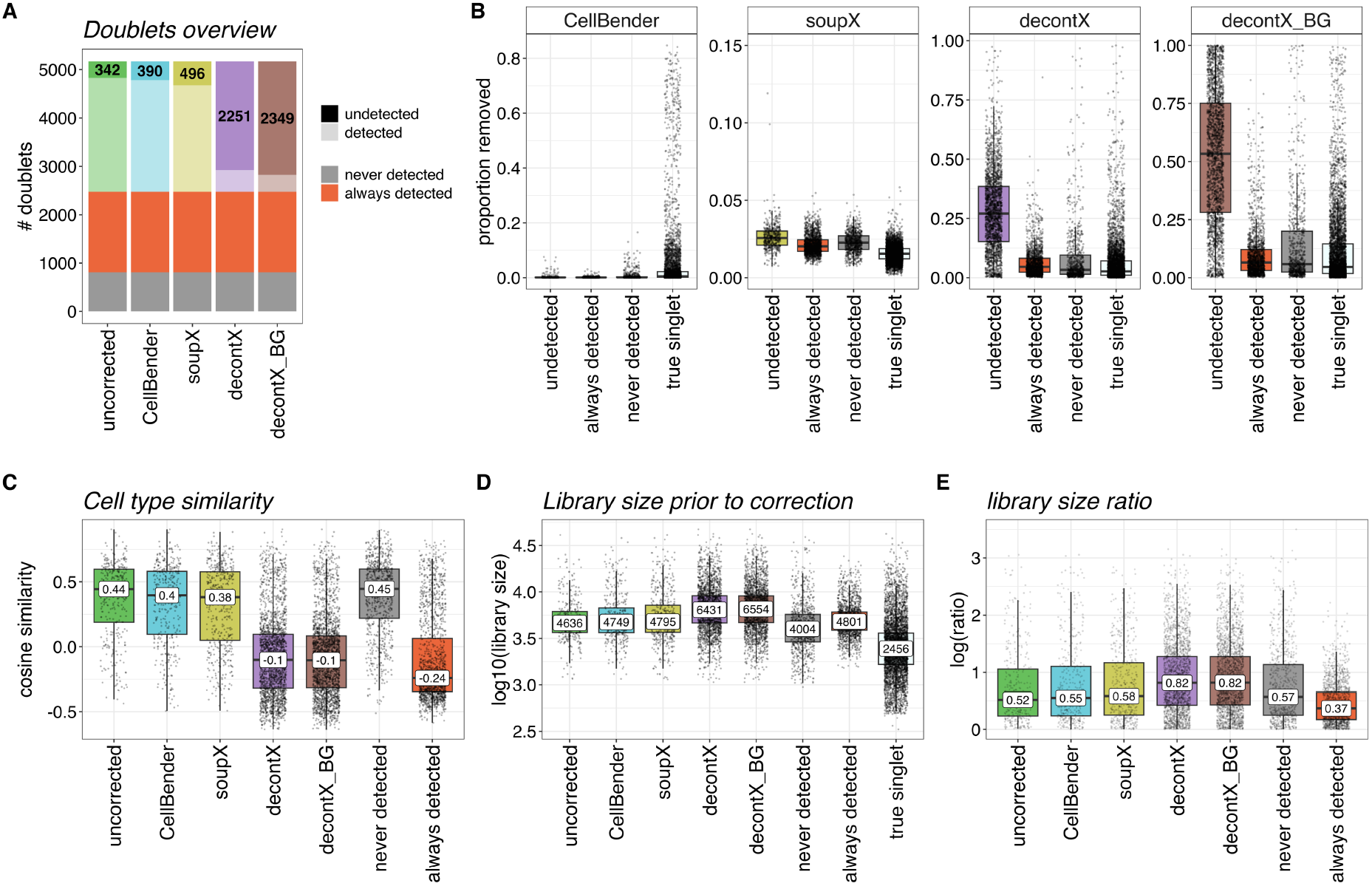
Engineered doublets reveal key features of undetected doublets after ambient RNA correction. **A.** Stacked barplot showing overview of doublet categories for all doublets (n=5,172) in all ambient RNA correction settings. The top portion (dark, numbered) represents the number of doublets that are not detected in that setting, which is referred to throughout the whole figure**. B.** Boxplots showing the proportion of counts removed by ambient RNA correction across sets of doublets or true singlets. Facets show the results for each ambient RNA correction tool. **C-E.** Boxplots characterising undetected doublets in each ambient RNA correction setting, as well as doublets that are always undetected, and doublets that are always detected. **C.** Boxplots showing the cosine similarity of two cells captured in a doublet. **D.** Boxplots showing the library size of doublets prior to ambient RNA correction. **E.** Boxplots showing the log library size ratio (larger cell/smaller cell). For all boxplots presented in this figure, the horizontal lines denote median values, boxes extend from the 25th to the 75th quantile, and whiskers extend to +/-1.5 inter-quantile range. Outliers are not highlighted, but all data points are shown individually. Labels refer to median values.

It has been previously reported that *decontX* removes a higher proportion of ambient RNA from doublets compared to singlets [35]. We therefore first investigate the proportion of counts removed in each droplet by each ambient RNA removal tool (Figure 4B). We find that overall *decontX* removes the most amount of signal compared to *CellBender* and *soupX*. In addition, while the proportion of counts removed by *CellBender* is highest in singlets, the proportion of counts removed by both *decontX* settings, and to some extent *soupX*, is increased in the doublets that are not detected after ambient RNA correction. This suggests that higher levels of ambient RNA removal likely interfere with doublet detection. A known limitation of most doublet detection tools is their inability to identify homotypic doublets [14,16,17] as well as their reduced sensitivity to doublets with smaller library size [14]. As expected, the two cells in doublets that are never detected show high cosine similarity (homotypic), while those always detected show low similarity (heterotypic) (Figure 4C). Interestingly, while doublets undetected in *soupX* or *CellBender* settings appear to have a high cosine similarity, those undetected in both *decontX* settings show low cosine similarity, suggesting transcriptional similarity is not the primary driver of misclassification following *decontX* correction. We next compare uncorrected library size across groups (Figure 4D). Doublets that are never detected have a smaller library size (mean = 4,004) than those always detected (mean = 4,801). In contrast, doublets that are not detected in all the ambient RNA settings have a larger library size than doublets that are never detected. This indicates that uncorrected library size is unlikely to explain misclassification after ambient RNA correction. Finally, we compare the library sizes of the two cells within a doublet by examining the logratio of the larger/smaller cell (Figure 4E). If the cells contribute the same amount of RNA this logratio will be 0 and larger values indicate more skewing between the cells. We find that all sets of undetected doublets show a higher library size ratio than the doublets which are always detected. This is particularly pronounced for doublets that are not detected in *decontX* settings. This suggests that while doublet calling is harder for droplets with uneven library size ratios in general [7], this effect is more pronounced after ambient RNA correction, especially with *decontX*.

Our analysis using engineered doublets provides valuable insight into general limitations of doublet detection. It demonstrates that the effectiveness of doublet calling is affected by the amount of RNA that is removed. It also allows us to disentangle why some doublets are more susceptible to increased ambient RNA correction, which ultimately leads to their misclassification as singlets. Specifically, we find that uneven cell contributions result in fewer detections after ambient correction using *decontX*.

### Doublet detection performs best on uncorrected or *CellBender* corrected data

We extend our analysis to include three additional datasets to assess how ambient RNA correction impacts doublet calling performance across a range of different biological contexts (Figure 5): the lung cancer dataset, the BAL dataset and the brain cortex dataset. Each dataset contains multiplexed samples, and we use detections across two samples as “true” doublets while acknowledging that within sample doublets are missed. The lung cancer dataset consists of non-small cell lung cancer dissociated tumour cells from 7 individual donors and the bronchoalveolar lavage (BAL) dataset [38] consists of mostly immune cells collected from 8 individual donors (4 healthy, 4 cystic fibrosis). Finally, as single-nuclei RNAseq data is known to be prone to ambient RNA contamination [32,33], we include the Brain Cortex dataset [39] which consists of nuclei from the brain cortex of 8 healthy, aging individuals.

**Figure 5.**
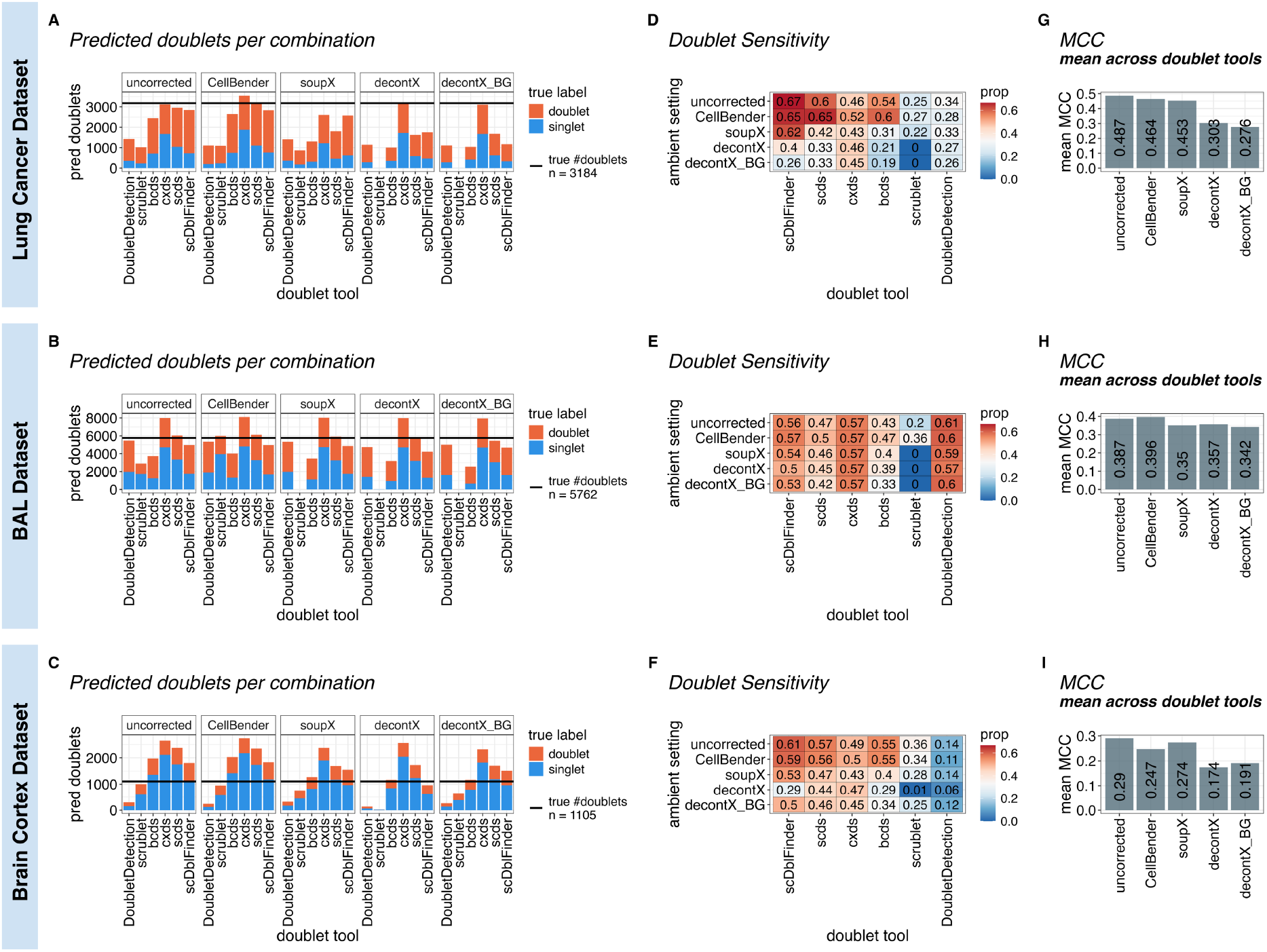
Doublet detection performs best on uncorrected or CellBender corrected data. **A-C**. Stacked barplots showing the number of predicted doublets for the lung cancer (**A**) BAL (**B**) and brain cortex dataset (**C**) The colour represents the proportion of true singlets (blue) and doublets (orange) of the predicted doublets. The solid line indicates the total number of true doublets. **D-F.** Heatmaps showing the proportion (prop) of true doublets detected for each combination of ambient RNA correction setting and doublet detection tool for the lung cancer (**D**) BAL (**E**) and brain cortex dataset (**F**). Colour scale represents proportion of true doublets detected within each dataset. **G-I**. Barplots showing mean MCC across all doublet detection tools for each ambient RNA correction setting for the lung cancer (**G**) BAL (**H**) and brain cortex dataset (**I**). Corresponding heatmap containing data for each doublet detection tool individually can be found in Supplementary Figure 4A-C.

For each dataset, we perform ambient RNA correction using four different approaches: two different settings for *decontX* (decontX and decontX_BG), *soupX* and *CellBender.* Doublets are identified using 6 different methods (*scDblFinder, scds, bcds, cxds, scrublet, DoubletDetection*). We argue that the erroneous classification of doublets as singlets is more problematic than the misclassification of singlets as doublets as it results in undetected doublets remaining in the dataset, potentially confounding downstream analyses. We therefore consider both the proportion of true doublets detected (doublet sensitivity) as well as MCC to evaluate doublet calling performance. A comprehensive list of additional performance metrics can be found in Extended Data Table 02.

Overall, we find that doublet calling performance in general is variable across different datasets (Figure 5A-C). For the Brain Cortex dataset in particular (Figure 5C), doublet calling performance is poor across all settings, due to high doublet false discovery rates (singlet predicted as doublet, blue proportion). The original authors observe that doublets in this dataset do not exhibit the usual higher library size compared to singlets [39], which likely interferes with some doublet detection algorithms, leading to reduced performance.

We find that different combinations of ambient RNA removal and doublet detection tools yield highly variable results and no combination performs best across all datasets. In general, we find that doublet sensitivity (proportion of true doublets detected) is highest in either the uncorrected or *CellBender* settings across all doublet detection tools (Figure 5D-F). While this is more pronounced in the Lung Cancer dataset (Figure 5D) and Brain Cortex (Figure 5F), the trend is consistent in all datasets. Importantly, across all doublet detection tools in all dataset, the uncorrected setting always matches or outperforms *soupX* and both *decontX* settings in doublet sensitivity. This suggests that, apart from *CellBender,* more true doublets are detected when doublet detection is performed on uncorrected data. Although doublet detection is impacted by ambient RNA correction for most doublet detection tools, we do note that *scrublet* appears to be particularly sensitive to ambient RNA correction performed with *soupX* and *decontX* resulting in none, or very few, doublets detected in some datasets (Figure 5D-F).

When looking at MCC (Figure 5G-I, Supplementary Figure 4A-C) we find that the mean across all doublet detection tools is highest in either uncorrected or *CellBender* settings. Therefore, in most cases, the benefit of increasing the detection of true doublets outweighs the slight loss in specificity. One notable exception is *soupX* in the Brain cortex dataset (Figure 5I, Supplementary Figure 4C). Here, *soupX* suffers from lower doublet sensitivity compared to *CellBender*, but its reduced false discovery rate rescues performance measured by MCC compared to *CellBender*. However, due to the overall poor performance of doublet calling in this dataset, it is unlikely to be exemplary of general trends across other datasets.

We further investigate the proportion of counts removed from undetected doublets in different ambient RNA correction settings (Supplementary Figure 4D-I). We selected *scDblFinder* to demonstrate these trends, as it detects the highest proportion of true doublets in most of the datasets. We find that *soupX* and *decontX* remove more RNA from doublets that are undetected after ambient RNA correction compared to doublets that are detected in all settings (Supplementary Figure 4G-I).

Despite variability in performance for doublet tools and across datasets, these results are consistent with the findings from engineered datasets (Figure 3). Specifically, doublet calling generates the most reliable results when performed on uncorrected or *CellBender* corrected data, and excessive RNA removal in undetected doublets by *soupX* and *decontX* indicates that these tools may remove doublet signals.

### A guide to the implementation order for ambient RNA and doublet detection tools

Ambient RNA correction undoubtedly improves analysis outcomes in many settings [30,32,33,40]. A recent study by Schroeder *et al* [40] succinctly summarises how to select an ambient RNA correction tool using a decision tree. A shortened version of this decision tree is presented in Figure 6A. The factors that most determine ambient RNA correction tool selection are data and compute availability. After selection of an appropriate tool, current workflows suggest that ambient RNA correction is performed prior to doublet detection.

**Figure 6.**
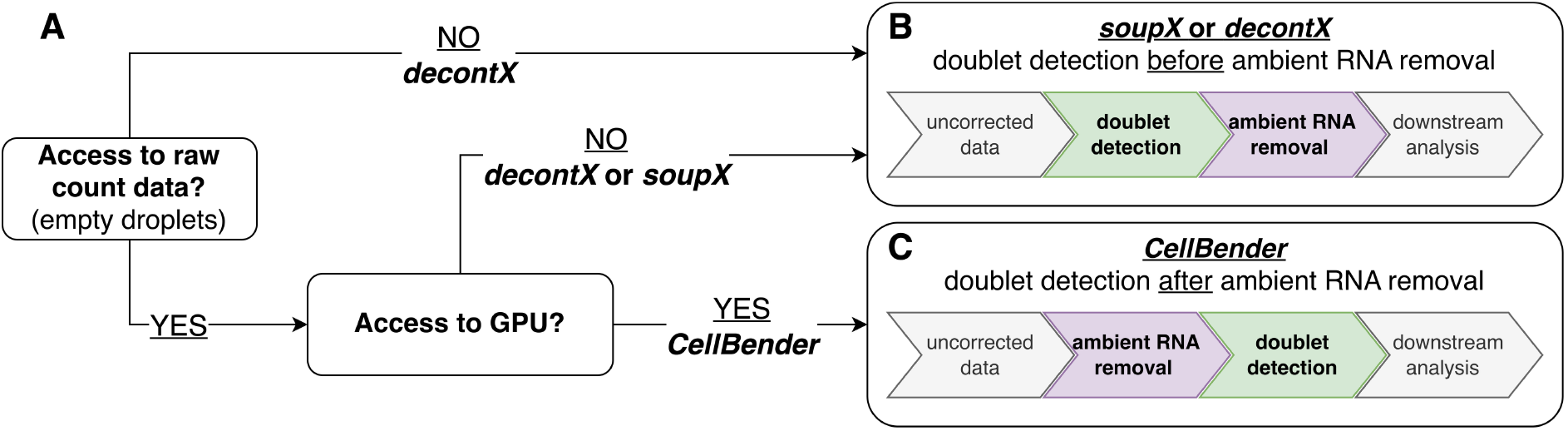
A guide to the implementation order for ambient RNA and doublet detection tools. **A.** Adapted and simplified decision tree presented by Schroeder et al. [40] to select ambient RNA correction tools based on data and compute availability. **B-C**. Extension of decision tree incorporating whether doublet detection should be performed before or after ambient RNA correction. For soupX and decontX, we recommend detecting doublets prior to ambient RNA correction (**B**), while we suggest running CellBender prior to doublet detection (**C**).

However, we find that for two of the three most used ambient RNA correction tools, doublet detection performance is reduced when compared to uncorrected data. This suggests that the sequence in which ambient RNA correction and doublet detection are performed requires reconsideration. We therefore extend the decision tree to also provide further information on whether doublet detection should be performed before or after ambient RNA correction (Figure 6).

Given the fact that doublet detection is improved in uncorrected data compared to *soupX* and *decontX* in most settings, and the ease at which the order of these tools can be rearranged, we recommend performing doublet detection prior to ambient RNA correction for *soupX* and *decontX* (Figure 6B). This mitigates the risk of ambient RNA correction removing doublet signal, and improves the performance of doublet detection. While ambient RNA correction using *soupX* and *decontX* can be applied at any point during the processing pipeline, *CellBender* can only be applied directly to raw, unfiltered count data. This is due to the fact that *CellBender* performs both cell-calling and ambient RNA correction, which naturally places *CellBender* upstream of any other form of processing. Given this requirement and the fact that applying *CellBender* mostly matches or improves doublet calling performance compared to uncorrected, running *CellBender* prior to doublet detection is relatively robust (Figure 6C).

## Discussion

In this study, we quantify the impact ambient RNA correction has on doublet detection performance using a variety of multiplexed datasets, with three commonly used ambient RNA correction tools (*soupX, decontX* and *CellBender*) and up to six different doublet detection tools (*scDblFinder, scds, cxds, bcds, scrublet, DoubletDetection*). We find that the highest doublet sensitivity is achieved when performing doublet detection on either uncorrected or *CellBender* corrected data. Based on our findings, we provide an easy-to-use guide to determine whether doublet detection should be run before or after ambient RNA correction for the three ambient RNA correction tools tested.

We show in a range of data sets that the proportion of predicted doublets decreases after ambient RNA correction is applied, indicating that ambient RNA correction may remove doublet signals. This effect appears to be more pronounced in biologically complex data, such as cancer data sets.

We next investigate doublets that remain undetected after ambient RNA correction. While multiplexed datasets using oligo/antibody-hashing or genetic diversity provide the necessary information to assess doublet detection performance, they do not offer insight into the composition of doublets. To address this, we introduce the concept of using 10x Flex data to generate engineered datasets, where both true doublet labels as well as doublet composition are known, and demonstrate their usefulness in evaluating doublet detection performance as well as characterising doublets that are not detected. Using engineered datasets, we quantify cell type similarity, which allows for distinction between homotypic and heterotypic doublets. We find that homotypic doublets are rarely detected in all settings. In *decontX* settings, however, most doublets that are not detected are heterotypic, but in these doublets the contribution of RNA from each cell is highly skewed. A much lower contribution of one cell to a droplet’s content may produce an expression pattern that resembles high levels of ambient RNA contamination, therefore leading to increased ambient RNA correction. Unequal contribution of cells to a doublet has previously been discussed in the context of optimising the simulation of doublets and presents an analytical challenge for doublet detection tools [7,19]. In agreement with this, our results show that doublets that are not detected in general exhibit uneven RNA contribution between their constituent cells. We next evaluate 30 combinations of doublet detection and ambient RNA correction tools using three multiplexed datasets that allow for doublet labelling between samples.

Experimentally labelled doublets have well-documented limitations as ground truth [16,19], including failure to detect same-sample doublets and dependence on demultiplexing accuracy. However, since we compare workflows with and without ambient RNA correction rather than evaluate tools, any ground truth imperfections affect all workflows equally and are unlikely to impact our overall conclusions. In addition, we use default or recommended parameters for tools rather than optimising parameters for individual tools. While we recognise that adjusting parameters for a given dataset may improve performance, evaluating tools under default settings remains meaningful given their widespread use in pipelines.

Overall, we find that doublet detection should be performed before ambient RNA correction, unless using *CellBender*. In particular, we find that doublet detection performance is strongly affected after ambient RNA correction using *decontX*. Importantly, *decontX* remains the only independently benchmarked tool with satisfactory ambient RNA correction performance that can operate without access to empty droplets [40]. This is a critical point for researchers re-analysing public data, in particular human data for which raw data often cannot be shared due to privacy reasons, or plate-based sequencing approaches which do not generate empty droplets [41]. Our results are therefore especially important for *decontX*-based workflows. A likely explanation for the reduced doublet detection performance after ambient RNA correction is that, with the exception of *CellBender* [34], the presence of doublets - particularly at high levels - is not explicitly modelled in ambient RNA correction tools. *decontX* models gene expression for each droplet as a mix of endogenous gene expression and contamination. This approach is inherently sensitive to doublets, which is discussed by the authors of the tool [35]*. soupX* derives an ambient profile from empty droplets and computes a global contamination rate, which, together with the total counts in the droplet, guides the removal of ambient RNA. As doublets tend to have a higher library size, this approach inflates the proportion of predicted ambient RNA in doublets. As a result, we find that doublets that are not detected after ambient RNA correction using *decontX* or *soupX* have an increased proportion of counts removed during ambient RNA correction.

Importantly, this is not a reflection on how well *soupX* and *decontX* perform at ambient RNA correction. Multiple studies have shown both tools perform well when used appropriately. However, as these tools are not designed to account for the presence of doublets, doublet detection should be performed on uncorrected data. In addition, removing doublets prior to ambient RNA detection is likely to ensure that the data is more compliant with the assumptions made by the tools thereby resulting in improved ambient RNA correction performance. In agreement with this, emerging ambient RNA correction tools [42] remove doublets prior to ambient RNA correction. Future methods incorporating a joint framework where doublets and ambient RNA correction are inferred simultaneously are likely to improve the detection of both.

## Conclusion

As scRNA-seq has become more accessible for large-scale studies, the volume of generated data necessitates automation of data processing. Consequently, scRNA-seq processing pipelines spanning different coding languages and ecosystems have emerged [36,37,43–45]. Many of these pipelines implement both ambient RNA correction and doublet detection. Here, we provide data-driven evidence to guide the order in which ambient RNA correction and doublet detection tools should be applied and, for the first time, demonstrate that doublet detection should be performed before ambient correction, unless using *CellBender*. These findings provide valuable evidence guiding informed decisions on how to implement these tools in pipelines, which will ultimately benefit the community.

## Resource availability

### Data Availability

The unfiltered and filtered Cell Ranger outputs for the 10x Flex tonsil dataset will be made available upon publication. All other datasets used in this study are publicly available via original publications [38,39,46] or websites [47–49].

### Code Availability

All the code used to generate the findings in this study can be found using this workflowR website (http://oshlacklab.com/doublet-ambient-paper/), or the corresponding GitHub repository (https://github.com/Oshlack/doublet-ambient-paper).

## Methods

Unless stated otherwise, all analysis was performed in R version 4.5.0 [50], Bioconductor version 3.21 [51] and package versions were managed through renv (version 1.1.4) [52]. We employ both SingleCellExperiment (version 1.30.1) [53] and Seurat (version 5.3.1) [54] throughout this study. Plots visualising results presented throughout this study were generated using ggplot2 (version 4.0.3)[55], patchwork (version 1.3.2) [56] and ggpubr (version 0.6.0) [57].

### Overview of datasets

Table 1 provides an overview of the datasets employed in this study. We use a selection of datasets across different biological conditions and sequencing technologies. Importantly, most datasets offer some form of multiplexing, either through surface lipids/proteins, genetic diversity, or barcode structure to provide ground truth doublet labels for doublet calling performance.

**Table 1.** Overview of datasets used in this study.

| Dataset name | Technology | Cells/<br>Nuclei | # samples | # donors | Tissue | Condition | Availability |
| --- | --- | --- | --- | --- | --- | --- | --- |
| PBMC dataset | 10X Flex v1 | cells | 16 | 1 | Blood | healthy | 10X Website: 128k Human PBMCs Stained with |
|  |  |  |  |  |  |  | TotalSeq™-C Human<br>Universal Cocktail<br>(Next GEM) |
| tonsil<br>dataset | 10X Flex v1 | cells | 16 | 16 | Tonsil | healthy | To be released upon<br>publication |
| ovarian<br>cancer<br>dataset | 10X Flex v1 | cells | 2 | 1 | Ovary | Cancer | 10X Website: Human<br>Ovarian Cancer FFPE<br>Single Cell Gene<br>Expression Flex (Next<br>GEM) |
| tonsil 10X 3'<br>v2 | Chromium 3'<br>(v2) | cells | 1 | 1 | Tonsil | healthy | E-MTAB-9005<br>(BCP002) |
| tonsil 10X 5'<br>v1 | Chromium 5'<br>(v1) + V(D)J | cells | 1 | 1 | Tonsil | healthy | E-MTAB-9005<br>(BCP003) |
| lung cancer<br>dataset | CellPlex +<br>Chromium 3'<br>(v3.1) | cells | 7 | 7 | Dissocia<br>ted<br>Tumour<br>Cells | Cancer | 10X Website: 20k<br>Mixture of NSCLC<br>DTCs from 7 donors, 3'<br>v3.1 (with intronic<br>reads) |
| BAL dataset | scHTO + ADT<br>+ Chromium<br>3' (v3.1) | cells | 8 | 8 | Bronchia<br>l lavage | Cystic<br>Fibrosis +<br>healthy | Zenodo<br>(https://doi.org/10.5281/zenodo.19381224)<br>batch 3 |
| Brain Cortex<br>dataset | ADT +<br>Chromium 3'<br>(v2) | nuclei | 8 | 8 | Brain<br>Cortex | healthy | docker container<br>(https://hub.docker.com/r/regevlab/demuxem<br>) , experiment4<br>1500nuc/ul |

For the PBMC [47] and ovarian cancer datasets [48], the filtered Cell Ranger output files were downloaded from the 10x website. For the lung cancer dataset [49], both the raw Cell Ranger output files as well as the demultiplexing outputs were downloaded. For the two non-flex tonsil samples, the Cell Ranger output files provided by the original authors were acquired [46]. For the BAL inflammation dataset [38], we downloaded the unfiltered Cell Ranger output for batch 3, as well as the processed object containing the *vireo* [10] derived demultiplexing results for that batch. We only considered one of the two captures for batch 3 (capture 3_1). For the brain cortex dataset [39], we downloaded the raw Cell Ranger output and ADT count matrix for experiment 4 (1500 nuc/ul) from the docker container provided by the original authors [39].

### 10x Flex tonsil dataset generation

For the tonsil dataset, samples were processed to single-cell suspensions, fixed within 1 hour of collection according to 10x Genomics Flex Protocol recommendations [58] and stored at -80°C until enough samples for one batch of 16 had been collected. Samples were thawed, barcoded, pooled (16 samples per pool), and processed according to 10x Genomics recommendations without modifications. For each capture, 128,000 cells were targeted, libraries were prepared using standard protocols, and sequencing was performed to achieve 10,000 reads/cell.

### Overview of tools

#### Doublet detection

In this study we test the effects of ambient RNA removal on doublet detection using six doublet detection tools, selected on the criteria of computational resources required, sensitivity to input parameters, popularity and accessibility.

*Solo* [22] is excluded due to its long runtimes and limited number of tutorials available to guide implementation. *Chord* [28] and *DoubletDecon* [23] are excluded due to their low popularity based on citations. *DoubletFinder* [16], despite being the most cited tool, is excluded for two reasons. Firstly, its doublet detection performance (measured by Matthews correlation coefficient) is considerably lower compared to other tools [14]. Secondly, our assessment framework uses the number of doublets predicted as a key performance metric for each method. However, with *DoubletFinder*, this is determined solely by the doublet proportion estimate provided as an input parameter, making direct comparison to other tools statistically unsuitable. The selected tools are briefly described below.

All doublet tools were implemented using modified run script templates provided by *Demuxafy* [14]. We removed droplets with very low counts (<200) or genes detected (<150) or particularly high proportion of mitochondrial reads (< 25% non-flex, <7% flex). As an exception, we did not remove droplets with high proportion of mitochondrial reads for the Ovarian Cancer dataset as certain tumour subpopulations in this dataset are known to exhibit high levels of mitochondrial reads. In addition, we removed droplets with unresolved genetic or hashtag origin from the Brain Cortex and BAL inflammation dataset. Doublet detection tools were then run on the resulting droplets, and both doublet prediction scores and labels were exported as csv files. Finally, doublet detection results from all tools were consolidated. Python packages were installed using micromamba (version 2.5.0) through miniforge3 with Python (version 3.14.3). To import Cell Ranger outputs for python-based tools, we adopted the functions provided by *Demuxafy*.

##### scds (bcds + cxds)[24]

*scds* identifies doublets using a combined approach: *cxds* identifies cells with co-expression patterns for mutually exclusive marker genes, while *bcds* generates artificial doublets, adds them to the original droplets and builds a gradient boosting classifier to identify doublets. Unlike other approaches which employ artificial doublets, *bcds* does not rely on dimensionality reduction and trains the classifier on log-transformed, normalised data instead. Finally, the framework also provides a *scds* as a hybrid approach which scales and combines *cxds* and *bcds* output scores. *Scds* is implemented in R and available through Bioconductor.

Doublet detection was performed using all three functions provided in the *scds* (version 1.24.0) package: both *cxds* and *bcds* were run using retRes=T and estNdbl=T parameters, followed by cxds_bcds_hybrid with estNdbl=T. We exported doublet calling results for all three approaches.

##### scDblFinder [7]

*scDblFinder* identifies doublets by generating artificial doublets and combining them with the original droplets, constructing a nearest neighbor network (kNN) and measuring the local neighbourhood density of artificial doublets for each droplet. Together with other predictors such as library size and marker co-expression, this neighbourhood score is then used to train a doublet classifier. *scDblFinder* is implemented in R and available through Bioconductor.

In preparation for doublet detection using *scDblFinder*, genes with fewer than 50 counts across all cells were removed and normalised gene expression values were calculated using the function logNormCounts from the *scuttle* (version 1.18.0) package. Dimensionality reduction was then performed using the runPCA followed by the runUMAP from the *scater* (version 1.36.0)[59] package. Finally, *scDblFinder* (version 1.20.0) was run using clusters = T and dbr.sd = 1. By setting dbr.sd = 1 we remove any assumptions made about expected doublet rates.

##### Scrublet [17]

*scrublet* identifies doublets by generating artificial doublets, combining them with the original droplets, constructing a nearest neighbor network (kNN) and assigning each droplet a doublet score based on the fraction of artificial doublets in the neighbourhood of each cell. Based on the distribution of the doublet scores for artificial doublets, a doublet threshold is determined to assign singlet and doublet labels to each droplet. *scrublet* is implemented in python and available through bioconda [60] or the scanpy [61] ecosystem.

*Scrublet* (version 0.3.2) was run using default parameters. In addition to the doublet calling results, we also export the distribution of simulated doublet scores and the position of the detected doublet threshold. We did not manually adjust the doublet detection threshold.

##### DoubletDetection[27]

*DoubletDetection* identifies doublets by generating artificial doublets, combining them with the original droplets and performing clustering. For all clusters, a hypergeometric test is performed to compute the artificial doublet enrichment in each cluster. Generation of artificial doublets, clustering and enrichment computations are repeated for a user-defined number of rounds. Finally, the average p-value across all rounds is used to determine doublets. Unlike most other doublet detection tools, *DoubletDetection* does not produce a doublet score, only binary labels. *DoubletDetection* is implemented in python and available through bioconda. *DoubletDetection* (version 4.3.0.post1) was run using the recommended parameter settings provided by the authors: n_iters=10, standard_scaling=True and pseudocount=0.1. As scanpy has deprecated louvain clustering, we implemented the recommended leiden clustering algorithm instead (clustering_algorithm=“leiden”, clustering_kwargs = {“flavor”: “igraph”}). In addition to the doublet calling results, we also export the convergence of doublet calls, as well as the number of doublets predicted at different threshold combinations for quality control. We did not manually adjust the voting threshold.

#### Ambient RNA correction

In this study, we selected the following three ambient RNA correction tools: *CellBender* [34], *decontX* [35] and *soupX* [31]. These tools were chosen as they generally perform well and are the most commonly used tools in the community. The selected tools are briefly described below.

##### CellBender [34]

*CellBender* is a deep generative model that distinguishes cell-containing and empty droplets, and performs ambient RNA correction. The content of each droplet is modelled as a mix of endogenous signal (cell expression) as well as background noise, such as ambient RNA, barcode swapping and capture efficiency. In addition to the denoised count integer count matrix, *CellBender* also provides information and guidance on model fitting and parameter inference. *CellBender* is implemented in python and available through bioconda.

*CellBender* (version 3.0.2) was run using the --cuda flag (cuda version 12.4.0) and the input parameters were specified for the individual datasets as reported in the table below. Expected cells and total droplets included were estimated based on kneeplots which were generated using the barcodeRanks function in the *DropletUtils* (version 1.28.0) package. Performance of *CellBender* was evaluated and input parameters adjusted as recommended by in-built reporting.

**Table 2.**
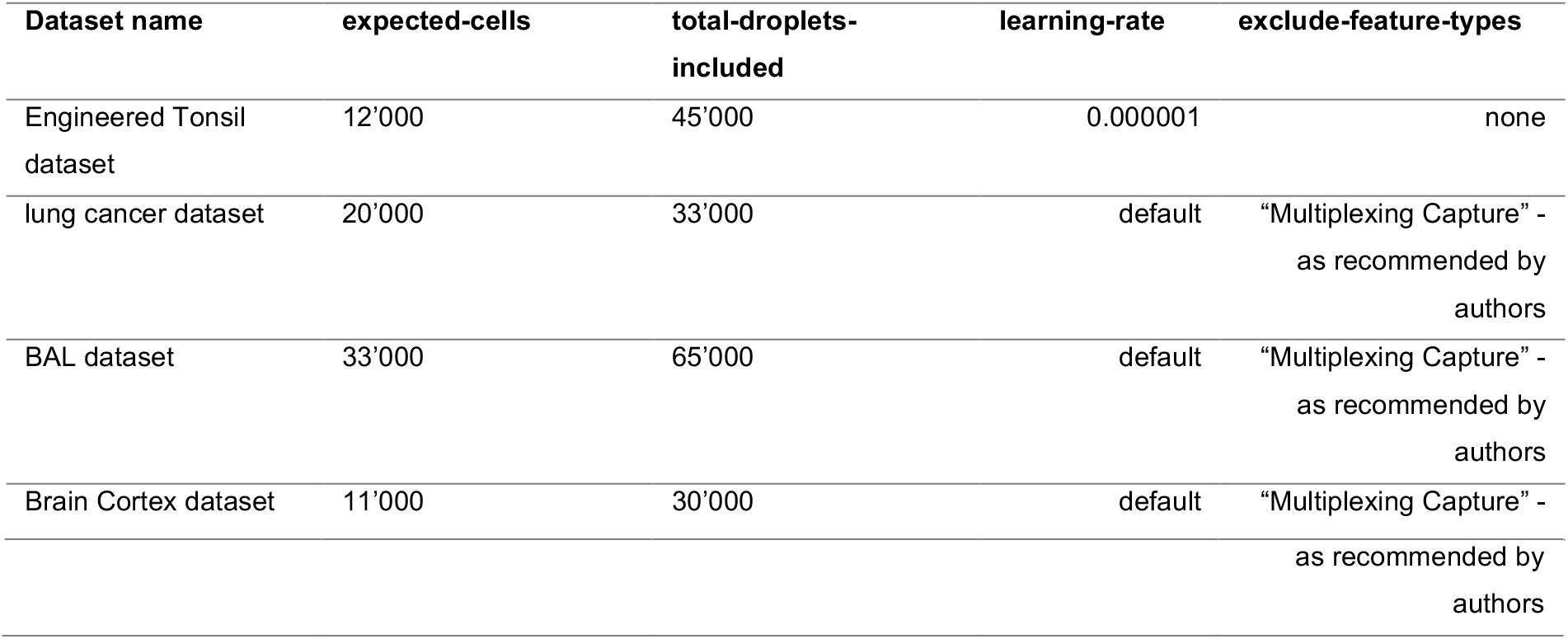
Overview of CellBender input parameters.

To make the resulting h5 files compatible with the *Seurat* function Read10X_h5, we converted them using the PyTables (version 3.7.0)[62] in python (version 3.7.12) to match the Cell Ranger output file format. For all downstream analysis, we only considered droplets that were detected as cells by Cell Ranger or barcodes provided by original authors, not *CellBender*. We exported the ambient RNA corrected counts matrix, the proportion of counts removed per cell and the proportion of genes detected lost per cell.

##### decontX [35]

*decontX* is a Bayesian approach to estimate and remove ambient RNA contamination in individual cells. The content in each droplet is treated as a mixture of two multimodal distributions- one from the endogenous expression of the cell, and the other from contamination. The model defines the contamination as a weighted combination of all other cell types present in the dataset. Cell type profiles are approximated by performing heuristic clustering. *decontX* can also be employed using empty droplets to empirically estimate the contamination distribution for each cell type from the background profiles instead. *decontX* is implemented in R and available through Bioconductor.

*decontX* (version 1.6.0) was run using default parameters. We employed both *decontX* settings, with and without background. For the background setting, we provided empty droplets (as identified by Cell Ranger or as provided by original authors) using the background parameter. We exported the ambient corrected counts matrix, the proportion of counts removed per cell and the proportion of genes detected lost per cell. The resulting corrected matrix was not converted to integers.

##### SoupX [31]

*SoupX* uses empty droplets to estimate and remove ambient RNA contamination. Gene expression abundance in empty droplets is utilised to derive a background contamination profile, followed by the estimation of a global contamination rate. This contamination rate can be computed by providing user-defined or automatically detected marker genes known to be absent in certain cell populations. Finally, the corrected count matrix is generated by subtracting the contamination profile, weighted by the estimated contamination rate and gene expression counts, from the original count matrix. *SoupX* is implemented in R and available through CRAN.

A SoupChannel was created using *SoupX* (version 1.6.2) with all non-empty droplets (as identified by Cell Ranger or as provided by original authors) as cells (toc parameter) and empty droplets as background (tod parameter). In further preparation for ambient RNA correction, dimensionality reduction was performed using the runPCA followed by the runUMAP from the *scater* (version 1.36.0) package. The resulting SingleCellExperiment was then converted to a Seurat object and louvain clustering using the FindNeighbors and FindClusters functions from the *Seurat* package was performed. FindNeighbors included 30 PCs, and FindClusters was run with resolution = 0.8. Clustering results and UMAP dimension reductions were added to the SoupChannel using the setClusters and setDR functions, respectively. We estimated the global contamination rate using the autoEstCont function with default parameters and finally performed ambient RNA correction using the adjustCounts function with default parameters. We exported the ambient corrected counts matrix, the proportion of counts removed per cell and the proportion of genes detected lost per cell. The resulting corrected matrix was not converted to integers.

### Evaluation of doublet detection performance

This study is not a benchmark for doublet detection tools. We are exclusively interested in quantifying how ambient RNA correction alters the performance of doublet detection. To this end, we implemented the following key performance metrics: The Matthews Correlation Coefficient (MCC) as well as the proportion of true singlets and true doublets detected.

#### Matthews Correlation Coefficient (MCC)

Doublet detection is a classification problem with very unbalanced group sizes, as singlets greatly outnumber doublets. For this reason, previous studies have used statistical metrics tailored to this, such as balanced accuracy, area under the precision recall curve (AUPRC) or MCC [14,19]. Critically, only MCC measures performance based on binary classification labels, whereas balanced accuracy and AUPRC operate on doublet scores. Most users do not rely on doublet scores but rather the binary classification label provided by the doublet detection tool to exclude or include droplets, therefore we use the MCC performance metric for this study. We computed the MCC for all tool combinations and datasets using the mcc function in the *yardstick* package (version 1.3.2) [63], using the factors singlet, doublet and Missing. Missing refers to droplets that no longer pass quality control filters after ambient RNA correction.

#### Additional doublet detection performance metrics

Traditional statistical metrics such as the MCC, while intuitive and informative, lack interpretability. To complement performance assessment using the MCC, we also quantify the proportion of true doublets and singlets detected. We mainly focus on the proportion of true doublets detected, which we refer to as doublet sensitivity. The following formulas outline how these metrics were computed:

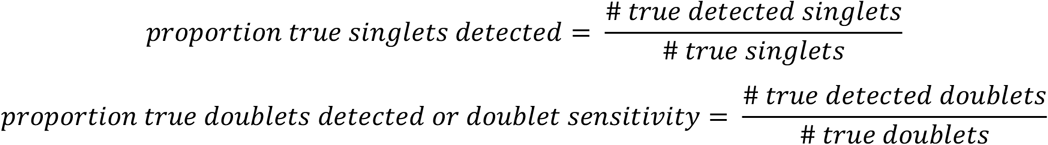

Finally, a range of additional performance metrics were computed using the confusionMatrix function in the *caret* package (version 7.0-1) [64] with mode = “everything”, as well as the pr_auc function in the *yardstick* package (version 1.3.2) [63].

### Generation of engineered dataset using 10x Flex

The following section outlines how we generate the engineered dataset (Figure 7A) and the corresponding empty droplets (Figure 7B) using the tonsil dataset. In this section, the combination of all cells carrying the same droplet barcode are referred to as *droplets*, while we refer to unique sample and droplet barcode combinations as *pseudo-droplets.* These are the output of the Cell Ranger pipeline for 10x Flex data, and are the equivalent of droplets in canonical single-cell experiments.

**Figure 7.**
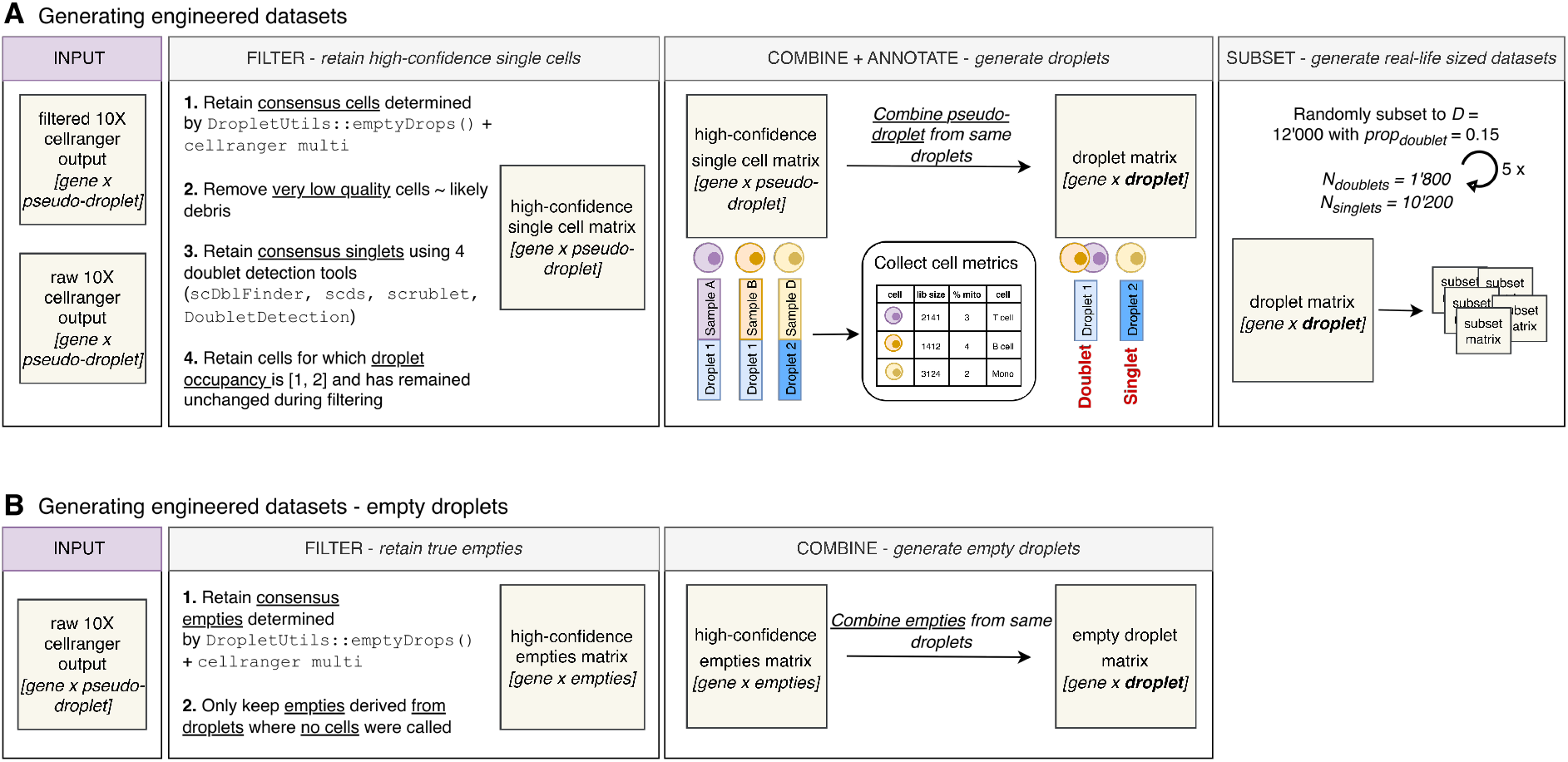
Generating engineered, ground truth datasets using 10x Flex data. **A.** Overview of how engineered datasets are generated from 10x Flex data**. B.** Overview of how the accompanying empty droplets are generated.

#### Generate engineered dataset

##### Filter - retain high-confidence singlets

We implemented various filtering steps to ensure that we are only considering high-confidence single cells as input for the engineered dataset. We only retain pseudo-droplets which were considered non-empty by both emptyDrops (*DropletUtils*, version 1.28.0) and cellranger multi (version 7.1.0)[2]. As the first 17 bases of the pseudo droplets barcode correspond to the droplet barcode, the droplet occupancy can be determined by quantifying how many non-empty pseudo-droplets carry the same droplet barcode. We refer to this as the original GEM occupancy. Next, we remove any pseudo-droplets that are of low quality (library size < 200, detected genes < 150 and mitochondrial reads > 7%) as they likely capture debris rather than real, intact cells. To ensure that engineered doublets never contain more than two cells, we identify and remove same-sample doublets. We employ four different doublet detection tools (*scDblFinder*, *scrublet*, *scds*, *DoubletDetection*) and determine the pseudo-droplets considered singlets by all four tools, which we term consensus singlets. By including multiple tools, we mitigate tool specific biases in singlet detection. Finally, we compute the post-filtering GEM occupancy and only pseudo droplets for which the original and post-filtering GEM occupancy remains the same are retained. Finally, we retain only pseudo-droplets whose GEM occupancy is either 1 (singlet) or 2 (engineered doublet).

##### Combine + Annotate -generate droplets

To generate engineered droplets, counts for pseudo droplets (now considered high-confident single cells) carrying the same droplet barcode are combined by summing the counts. In addition, we collect metadata information for individual cells in each engineered droplet. To quantify transcriptional similarity between cells within a doublet, we also compute the cosine similarity between cells using the cos_matrix function in the *coop* (version 0.6-3) package [65].

##### Subset -generate real-life sized datasets

To test our workflow in real-world lab conditions, we subset the full counts matrix by random sampling using the base R sample function, so that the subset includes 1,800 doublets and 10,200 singlets, resulting in a total engineered dataset size of 12,000 engineered droplets with a doublet proportion of 0.15. We repeat this process 5 times to generate multiple batches of engineered datasets. The final engineered datasets are exported in Cell Ranger format using the write10xCounts function in the *DropletUtils* (version 1.28.0) package with version = “3”.

#### Generate empty droplets background

As most ambient RNA correction tools require empty droplets to generate a background profile, we also generate an appropriate empty droplet background for the engineered datasets. Empty pseudo-droplets, which we refer to as *empties,* are obtained from the pseudo-droplets called empty by both emptyDrops (*DropletUtils*, version 1.28.0) and cellranger multi. We further restrict ourselves to pseudo-droplets that are captured in droplets in which no cells were detected. Empty droplets are obtained by summing the counts of empties which carry the same droplet barcode. As *CellBender* requires unfiltered data, we combined the empty and cell-containing droplets.

### Workflow in 10x Flex datasets

We remove ambient RNA contamination from all 10x Flex datasets (tonsil dataset, PBMC dataset, ovarian cancer dataset) using *decontX* without background as described above. We perform doublet calling with *scDblFinder* as described above, except that we use the samples parameter, which runs *scDblFinder* on each sample individually.

For the tonsil dataset, we perform louvain clustering using the FindNeighbors and FindClusters functions from the *Seurat* (version 5.3.1) package. FindNeighbors included 30 PCs, and FindClusters was run with resolution = 0.1 and algorithm = 3. Clusters were annotated manually by visualising gene expression for *CD79A* (B cells) and *CD3E* (T cells). UMAP visualisations were created using the plotUMAP function from the *scater* package (version 1.36.0).

### Workflow in multiplexed datasets

Ambient RNA contamination was removed from all datasets (lung cancer dataset, BAL dataset, brain cortex dataset) using all ambient RNA correction and doublet detection tools as described above. We evaluated the quality of hashtagging for the lung cancer and BAL dataset as previously described [12] to confirm that these datasets would be able to provide trustworthy ground truth labels. For all datasets, ground truth was defined as droplets which both pass quality control filters using the raw data (as described above), and for which demultiplexing results were available. The following table summarises what was considered the ground truth doublet/singlet labels for each dataset.

**Table 3.**
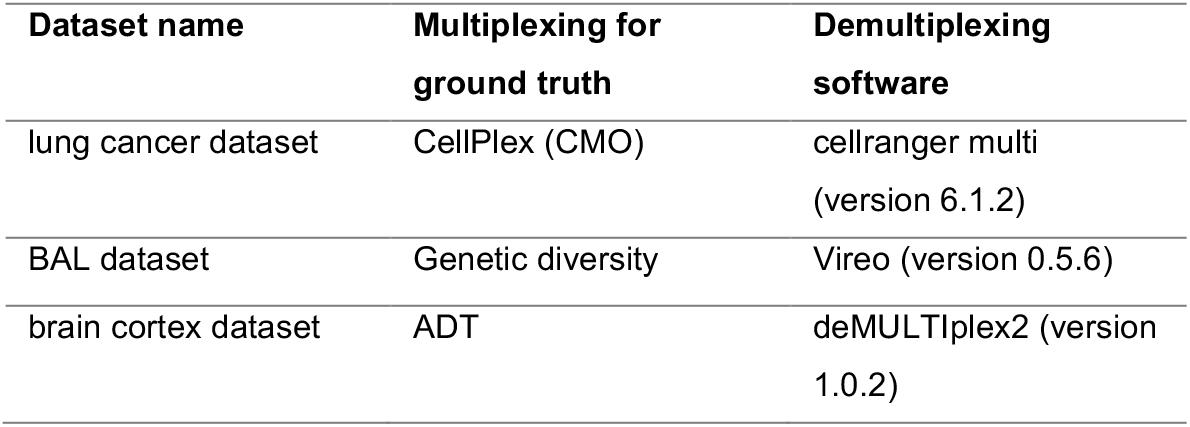
Experimental doublet/singlet labels for multiplexed datasets.

| Dataset name | Multiplexing for ground truth | Demultiplexing software |
| --- | --- | --- |
| lung cancer dataset | CellPlex (CMO) | cellranger multi (version 6.1.2) |
| BAL dataset | Genetic diversity | Vireo (version 0.5.6) |
| brain cortex dataset | ADT | deMULTiplex2 (version 1.0.2) |

For both the lung cancer as well as the BAL dataset, the demultiplexing results provided by the original authors were used. For the brain cortex dataset, demultiplexing results were generated using *deMULTIplex2* (version 1.0.2) [66] with default parameters.

## Supporting information

Extended Data Table 02: All performance metrics for Figure 05

Extended Data Table 01: All performance metrics for Figure 03

## Declarations

### Use of large language models

Large language model (LLM) Claude (Anthropic, Sonnet 4.6) was used to optimise code used for visualisation purposes. All data analysis, writing and interpretation of results were performed without LLM assistance.

## Supplementary Data

## Supplementary Data

**Supplementary Figure 1.**
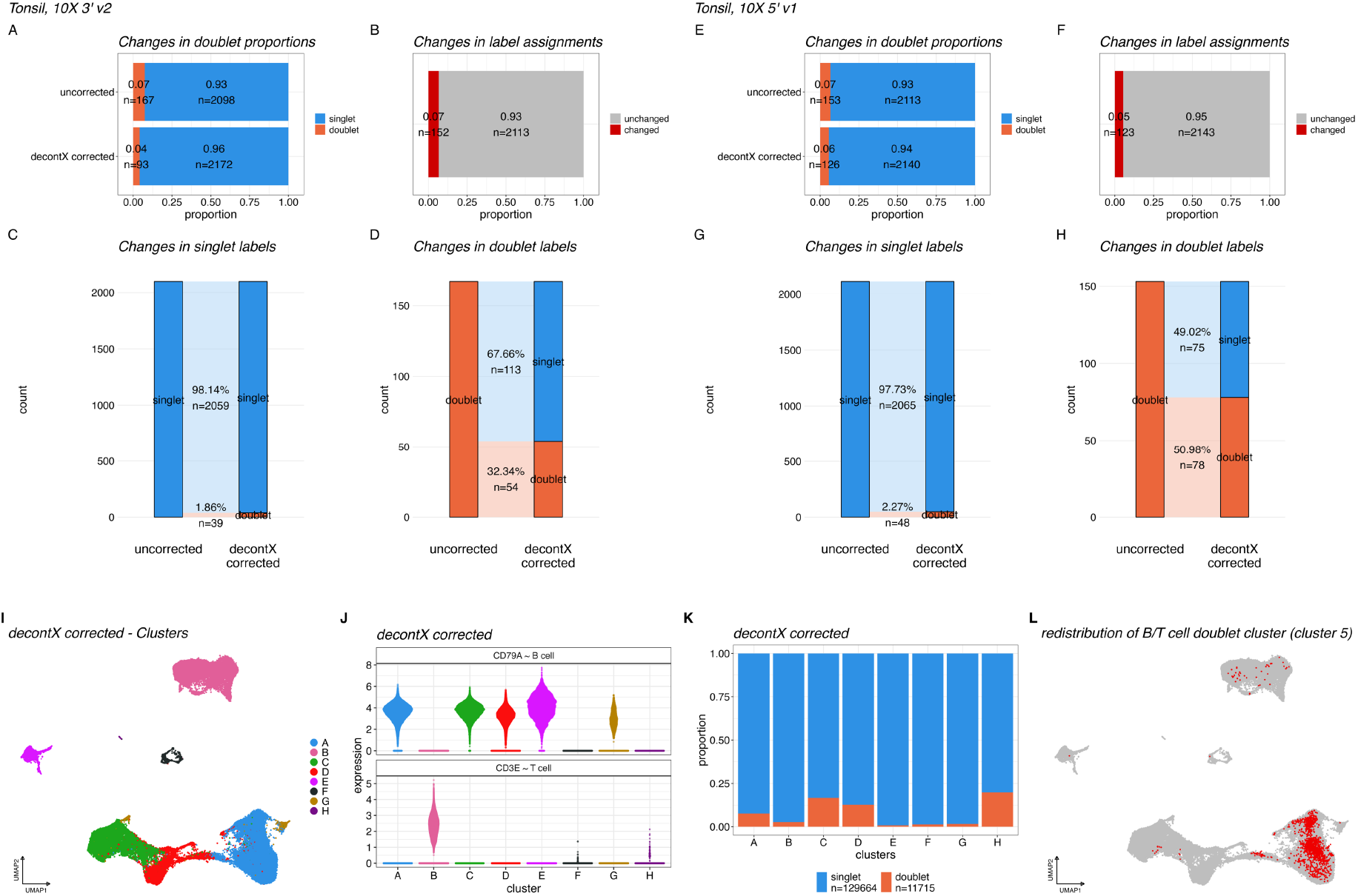
Performing ambient RNA correction alters doublet detection outcomes in non-flex datasets. (**A-D**) show results for tonsil cell data generated using the 10x 3’ v2 and (**E-F**) using 10x 5’ v1 technology **A+E**. Stacked barplots showing the proportion of doublets and singlets in the uncorrected (top) and decontX corrected (bottom) data in the 10x 3’ v2 (**A**) and 10x 5’ v1 (**E**) dataset. **B+F**. Stacked barplots showing the proportion of cells whose predicted labels change after ambient RNA correction in the 10x 3’ v2 (**B**) and 10x 5’ v1 (**F**) dataset. **C-D**. River plots showing how predicted labels change for singlets (**C**) and doublets (**D**) in the 10x 3’ v2 dataset. **G-H**. River plots showing how predicted labels change for singlets (**G**) and doublets (**H**) in the 10x 5’ v1 dataset. **I.** UMAP visualisation showing clusters in the decontX corrected tonsil dataset. **J.** Violin plot showing gene expression for CD79A (B cell marker) and CD3E (T cell marker) for the clusters in the decontX corrected dataset. **K.** Stacked barplots showing the proportion of doublets and singlets for each cluster in the decontX corrected dataset. **L**. UMAP visualisation highlighting the re-distribution of for B/T cell doublet cluster (cluster 5) cells (black, n=1911) in the decontX corrected dataset.

**Supplementary Figure 2.**
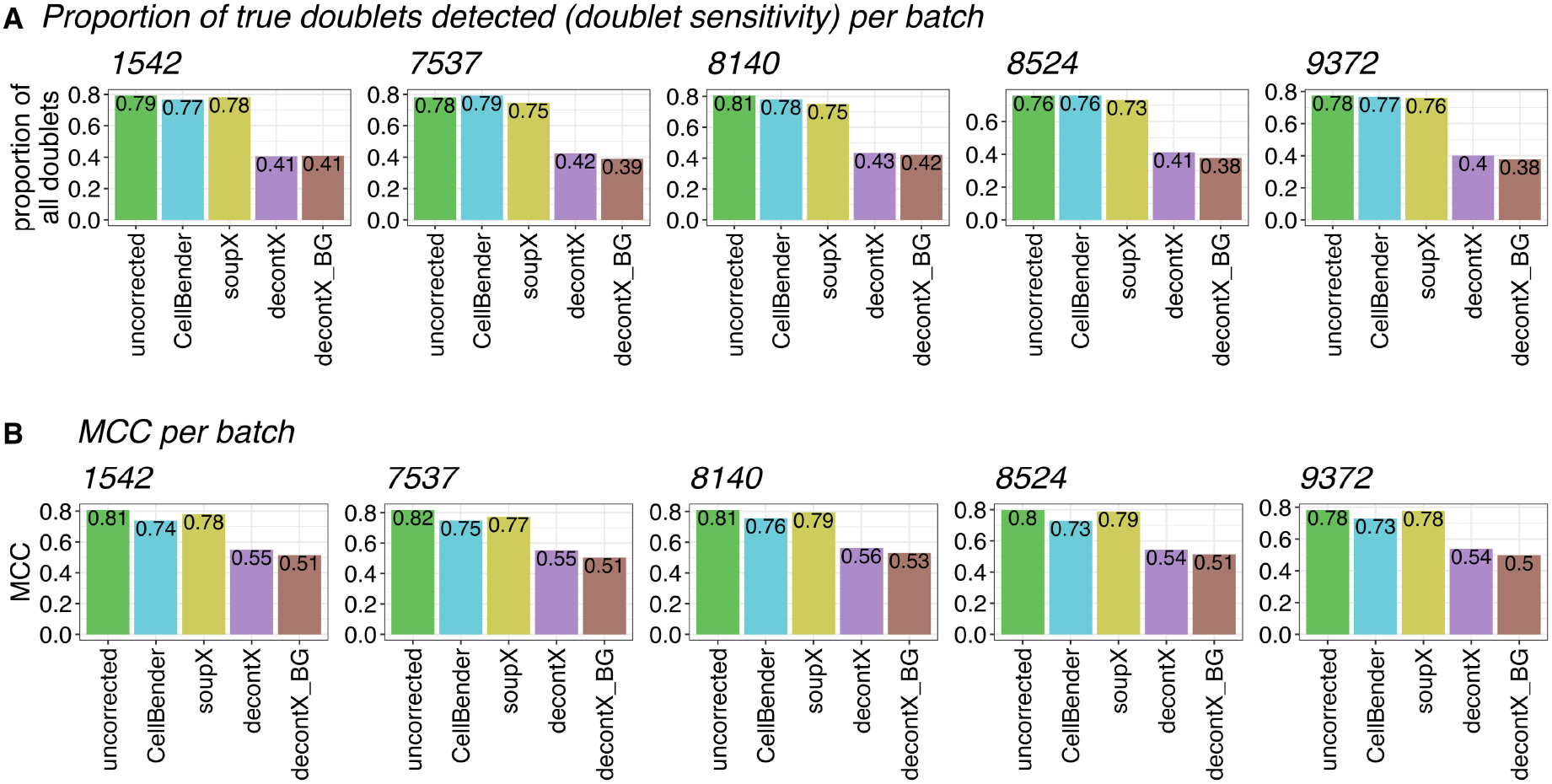
Doublet detection performance for each batch of the engineered tonsil dataset. **A-B.** Barplots showing the proportion of true doublets detected (**A**) or MCC (**B**) for each ambient RNA correction setting. Each plot shows results for one batch.

**Supplementary Figure 3.**
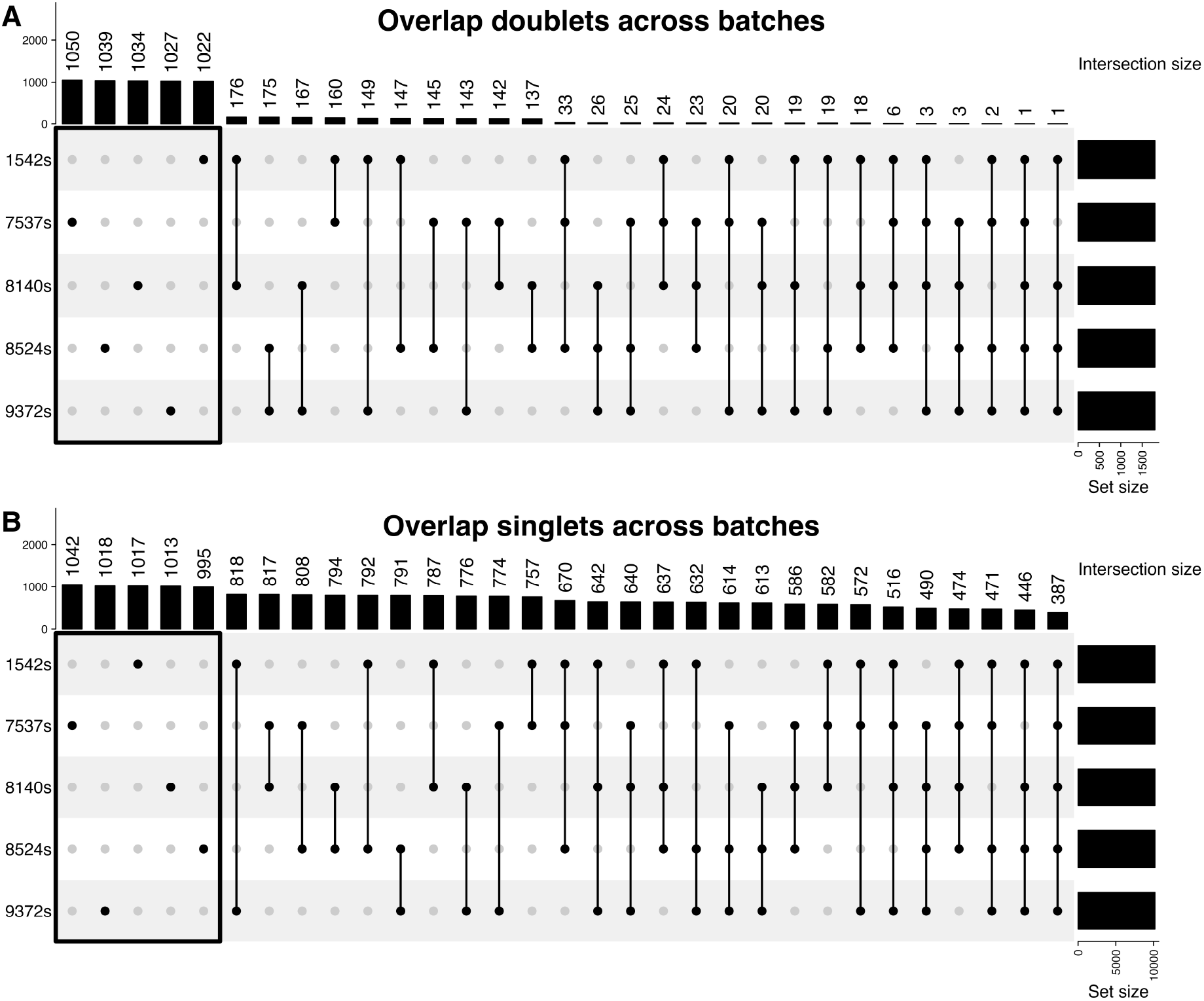
Unique doublets and singlets across engineered dataset batches. **A-B.** Upset plots showing intersections of true doublets (**A**) and singlets (**B**) across all batches. The top barplots display the size of each combination. For the analysis presented in Figure 4, we select doublets and singlets that are only present in one batch (highlighted with box).

**Supplementary Figure 4.**
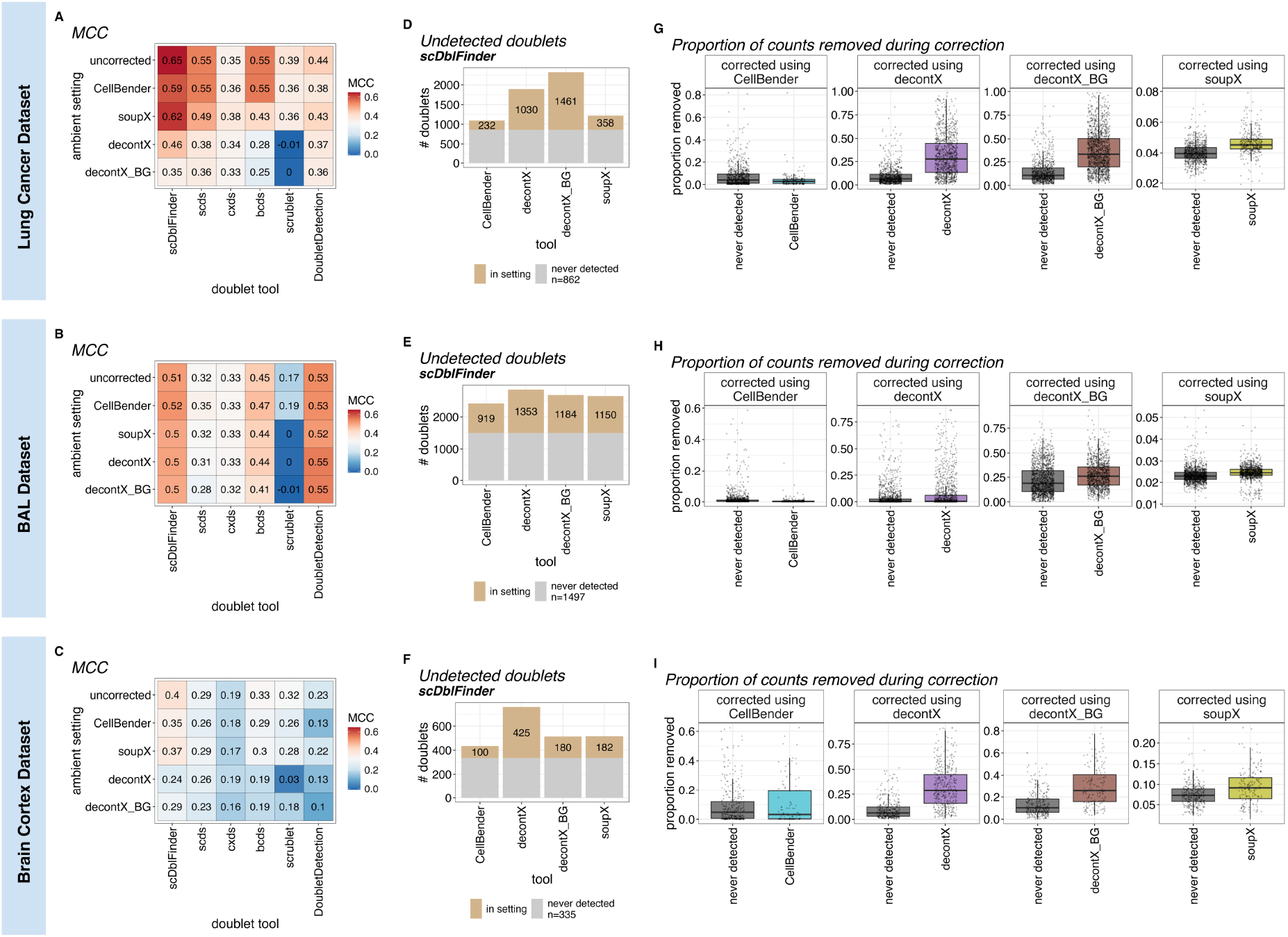
Doublet detection performs best on uncorrected or CellBender corrected data. **A-C.** Heatmaps showing MCC for each combination of ambient RNA correction setting and doublet detection tool for the lung cancer (**A**) BAL (**B**) and brain cortex dataset (**C**). Colour scale represents MCC. **D-F**. Stacked barplot showing the number of undetected doublets for each ambient RNA correction tool, split by the proportion that is shared across all settings (grey, always undetected) and the proportion that corresponds to undetected doublets in that setting (tan). **G-I.** Boxplots showing the proportion of counts removed by ambient RNA correction for doublets that are never detected (grey) and doublet undetected in the corresponding ambient RNA correction setting. Each facet shows the results for an ambient RNA correction tool. For all boxplots presented in this figure, the horizontal lines denote median values, boxes extend from the 25th to the 75th quantile, and whiskers extend to +-1.5 inter-quantile range. Outliers are not highlighted, but all data points are shown individually.

